## Supplemental data complete for "Retroviral integration into nucleosomes through DNA looping and sliding along the histone octamer"

#### ^2^Chromatin structure and mobile DNA Laboratory, The Francis Crick Institute, NW1 1AT, London, UK

^3^Single Molecule Imaging Laboratory, MRC London Institute for Medical Science, W12 0NN, London, UK

^4^Structural Biology Science Technology Platform, The Francis Crick Institute, NW1 1AT, London, UK.

^5^Molecular Virology, Department of Medicine, Imperial College London, W12 0NN, London, UK

^%^Present address: Wellcome Centre for Cell Biology, University of Edinburgh, EH9 3JR Edinburgh, UK

^#^Present address: NeCEN, University of Leiden, 2333CC, Leiden, The Netherlands

### ^&^Present address: Faculty of Biological Sciences, LS2 9JT, Leeds, UK

^§^Equal contribution.

*Correspondence:.

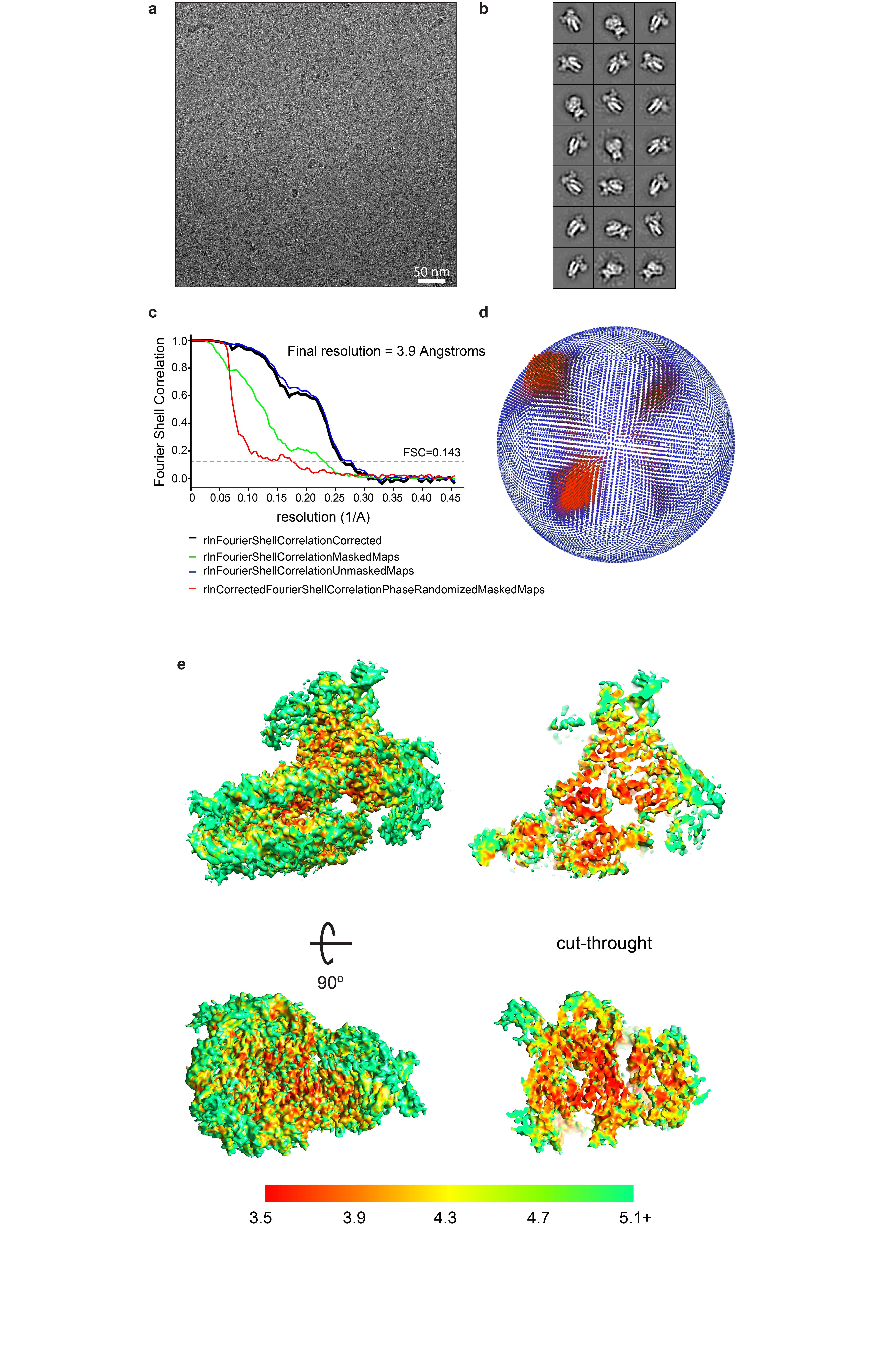

**Supplementary Figure 1** Cryo-EM analysis of Intasome-D02 NCP. **(a)** Representative micrograph. **(b)** Representative 2D averages. **(c)** Gold standard FSC curve for the refined cryo-EM map. **(d)** Euler angle distribution plot for all particles included in the final map. Bar length and colour (blue low, red high) correspond to number of particles contributing to each view. **(e)** Cryo-EM map coloured according to local resolution estimated with ResMap.

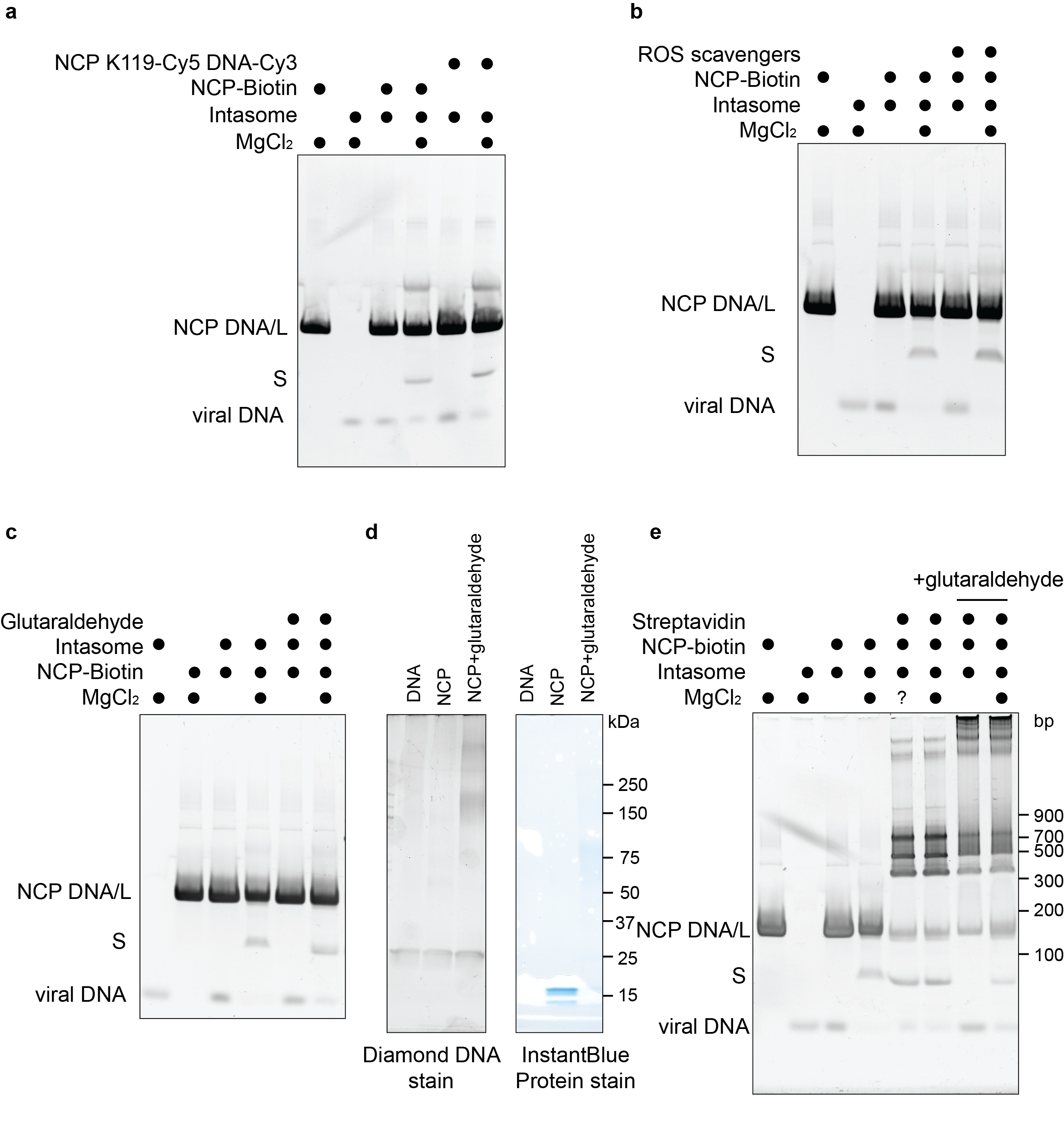
**Supplementary Figure 2** Retroviral integration into various nucleosome derivatives. **(a)** Integration assay shows magnesium dependence of viral strand transfer. Strand transfer product S and L labeled. Assay performed with D02-NCP used for structure determination and fluorophore labelled nucleosomes used in single-molecule FRET experiments. **(b)** PFV Integration assay shows that ROS scavengers used in single-molecule FRET experiments do not affect the strand transfer reaction. **(c)** Glutaraldehyde cross-linking of D02 NCP does not prevent integration. **(d)** SDS-PAGE gels stained with Diamond nucleic acid stain (left) or InstantBlue protein stain (right). Glutaraldhyde cross-inked sample migrates as lower electrophoretic-mobility smear. **(e)** Efficient integration occurs with NCP-D02-biotin complexes pre-assembled with streptavidin, and cross-linked with glutaraldehyde.

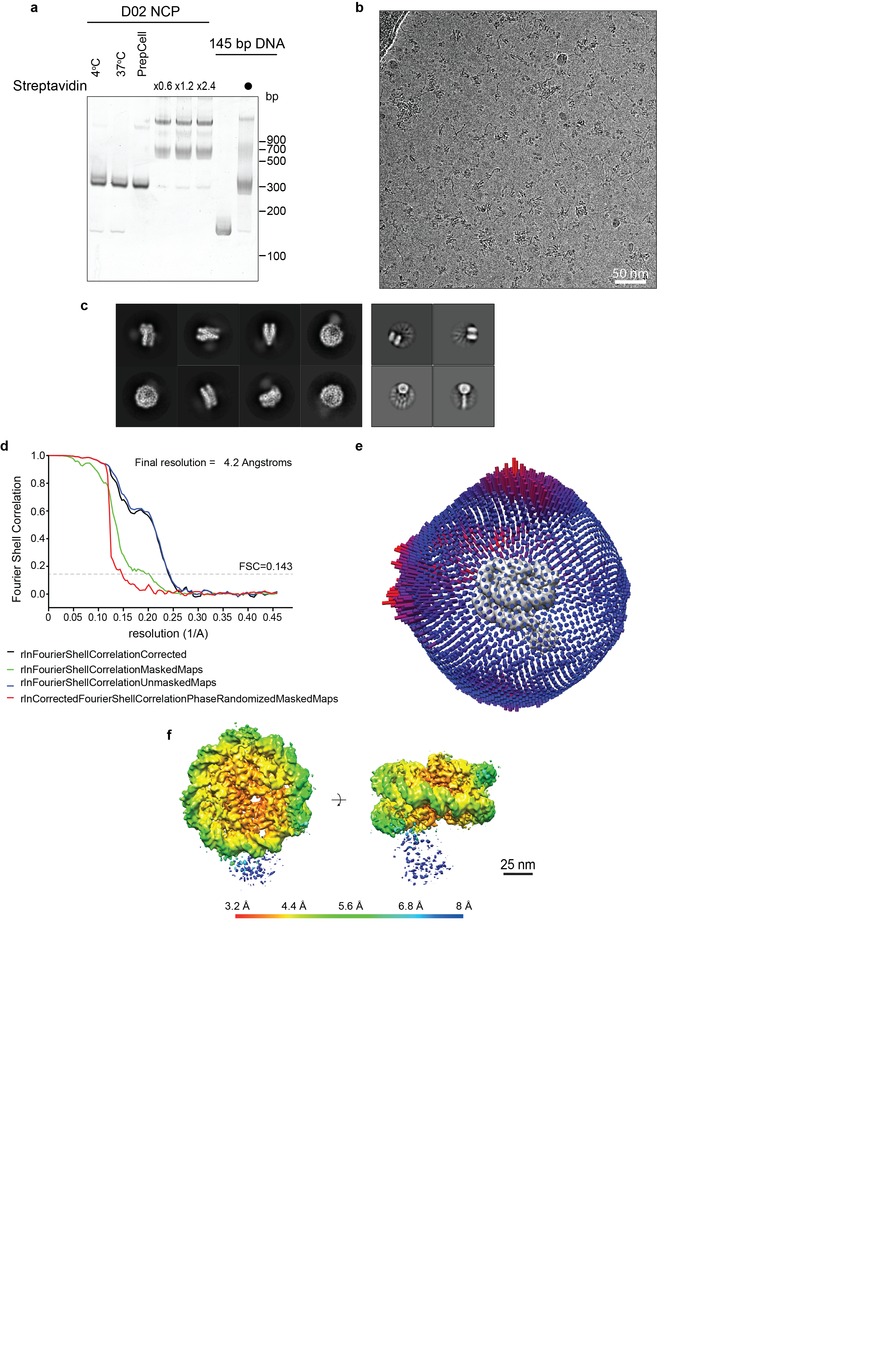

**Supplementary Figure 3** D02 NCP-streptavidin preparation and cryo-EM characterisation. **(a)** Native gel electrophoresis of biotin-D02 wrapped NCP during purification and interaction with streptavidin. **(b)** Representative cryo-EM micrograph of NCP-D02-streptavidin. **(c)** Selected 2D averages of D02 NCP-streptavidin (left) and isolated or DNA-bound streptavidin (right). **(d)** Gold standard FSC curve for the final structure. (**e**) Euler angle distribution plot for all particles included in the final map. Bar length and color (blue low, red high) correspond to number of particles contributing to each view. (**f**) NCP-D02-streptavidin coloured according to local resolution estimated in RELION. Streptavidin density is fragmented at the displayed threshold, due to the flexible tether with DNA.

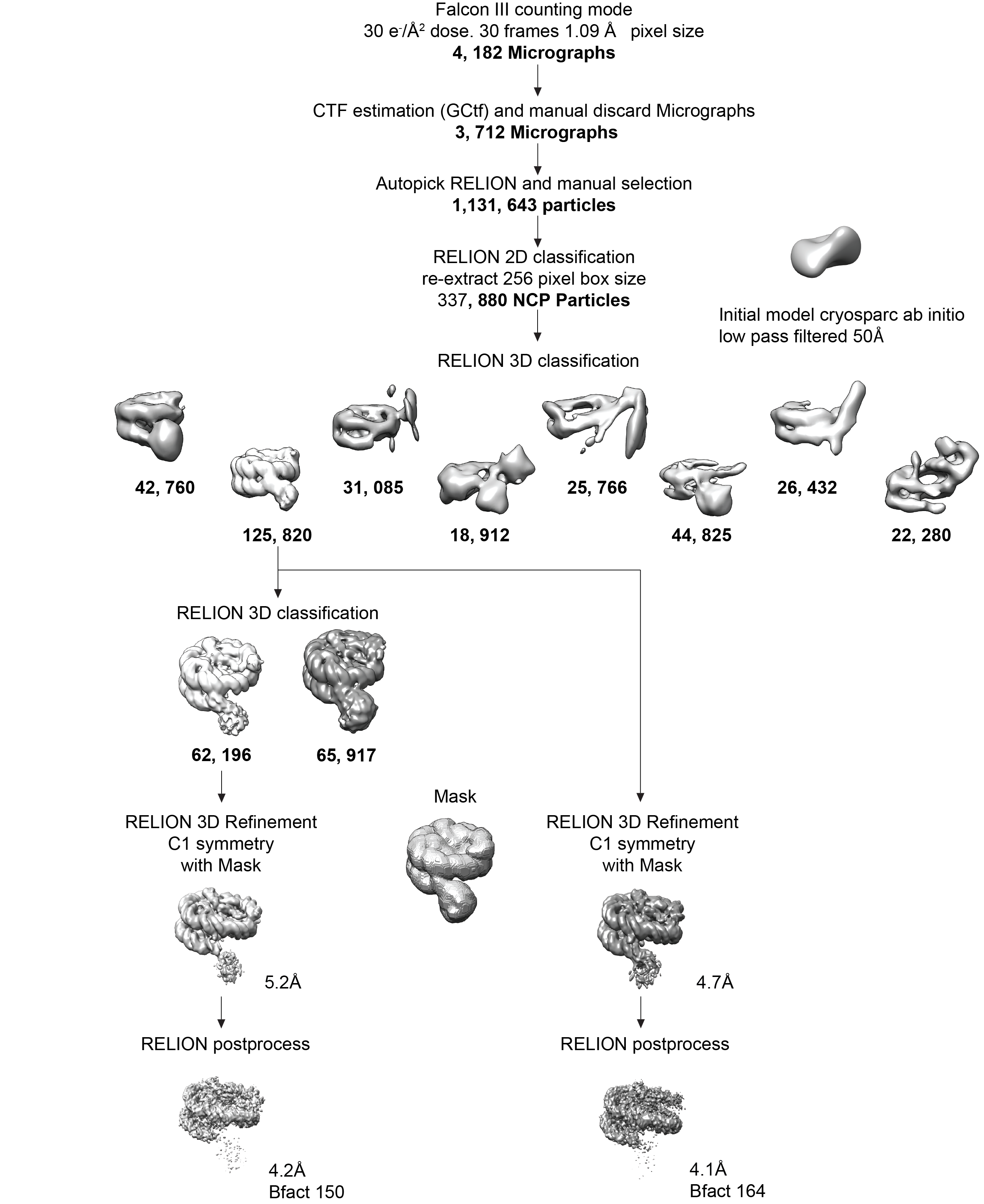

**Supplementary Figure 4** Overview of cryo-EM single-particle reconstruction for D02 NCP-streptavidin.

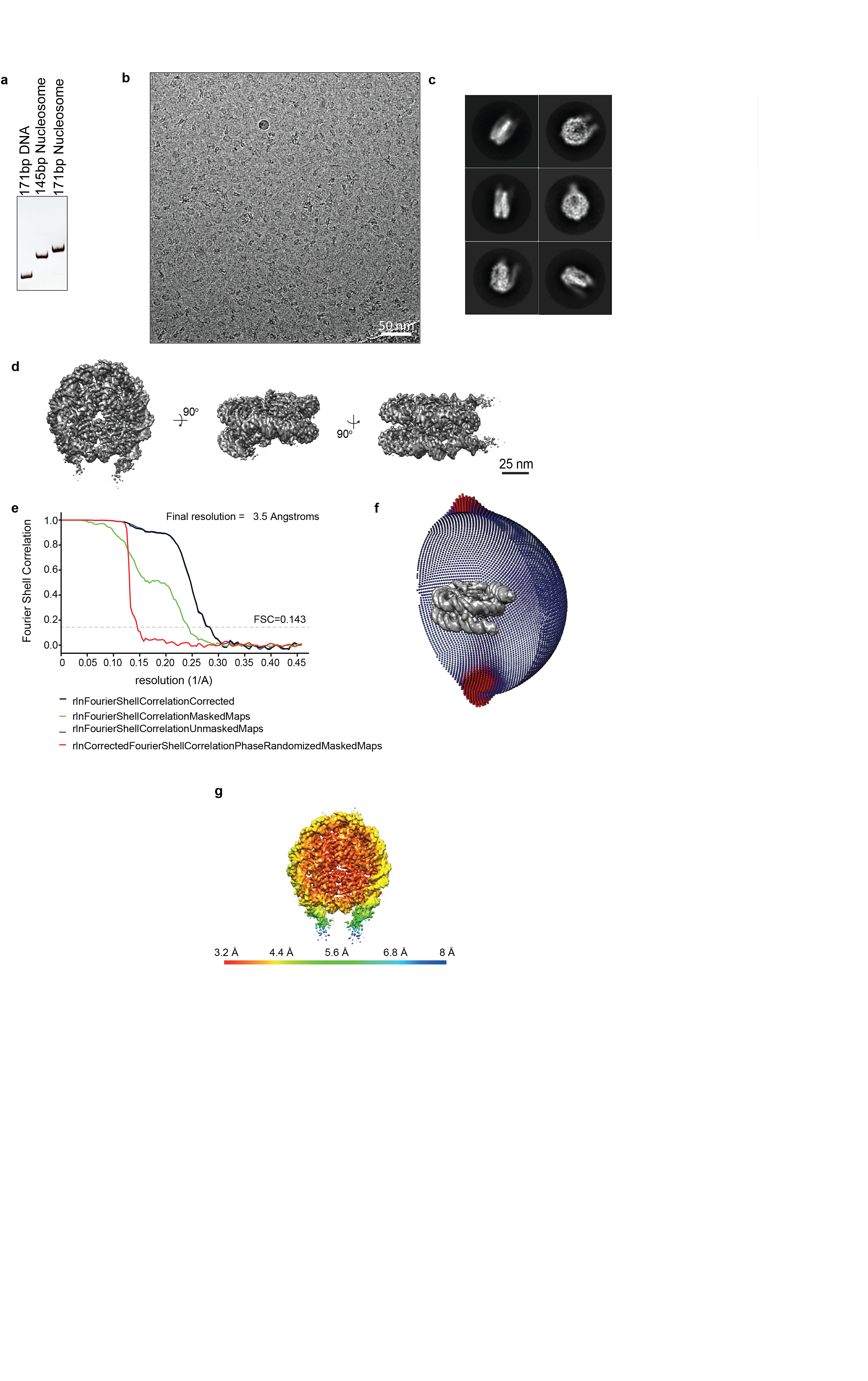

**Supplementary Figure 5** Electron microscopy of Widom 601 NCP. **(A)** Native gel electrophoresis of Widom 601 DNA isolated, and wrapped in a nucleosomes with either 145 base pair or 171 base pair-long DNA. **(B)** Representative cryo micrograph of nucleosome reconstituted with 171bp of Widom 601 DNA. **(C)** Selected representative 2D class averages. **(D)** Surface rendering of the NCP 171bp complex, shown in three views rotated about their axis. **(E)** Gold standard FSC curve for the refined cryo-EM map. **(F)** Euler angle distribution plot of all particles used for the final map (C2 symmetry imposed. Bar length and colour (blue low, red high) corresponds to number of particle images contributed to each view. **(G)** 171-base pair Widom 601 nucleosome, coloured according to local resolution estimated in RELION. Density for the DNA projecting away from the NCP core is visibly weaker.

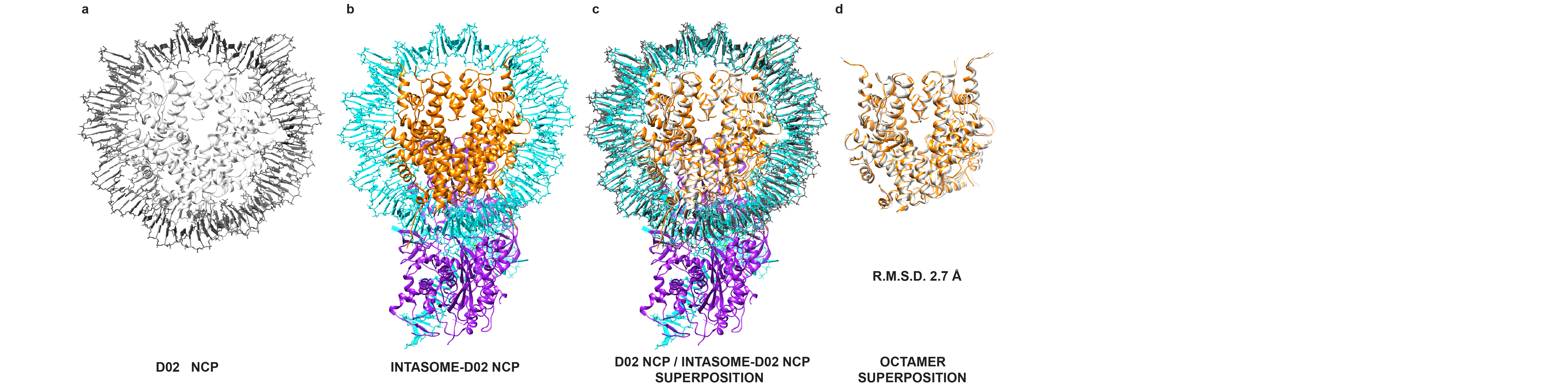
**Supplementary Figure 6** Retroviral integration causes octamer compaction. **(a)** Intasome-free D02 NCP alone, **(b)** Intasome-NCP structure, **(c)** the two structures with octamers superposed and **(d)** isolated octamers. Superposition causes slight octamer compaction (RMSD 2.7 Å).

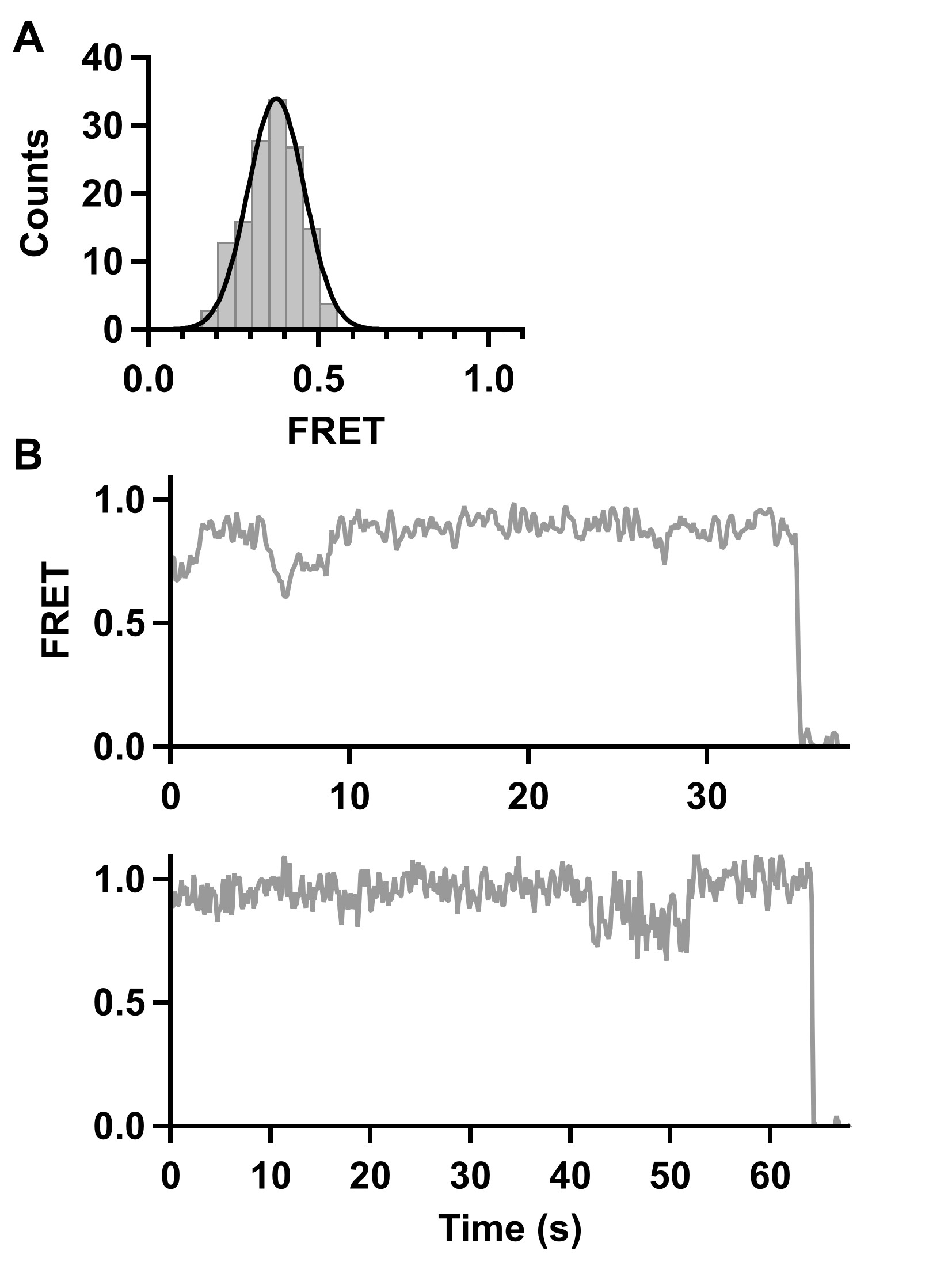
**Supplementary Figure 7 Single molecule FRET of D02 nucleosome core particles (A)** Single-molecule FRET histogram of distal-only labeled nucleosomes (N = 140). Black line is fit to a Gaussian distribution. **(B)** Representative dynamic single-molecule FRET time trajectories of proximal-only fluorescent nucleosomes. Data were collected at 100 ms/frame and smoothed with a 3-point moving average.
